# Dynamic neurotransmitter specific transcription factor expression profiles during *Drosophila* development

**DOI:** 10.1101/830315

**Authors:** Alicia Estacio-Gómez, Amira Hassan, Emma Walmsley, Lily Wong Le, Tony D. Southall

## Abstract

The remarkable diversity of neurons in the nervous system is generated during development, when properties such as cell morphology, receptor profiles and neurotransmitter identities are specified. In order to gain a greater understanding of neurotransmitter specification we profiled the transcription state of cholinergic, GABAergic and glutamatergic neurons *in vivo* at three developmental time points. We identified 86 differentially expressed transcription factors that are uniquely enriched, or uniquely depleted, in a specific neurotransmitter type. Some transcription factors show a similar profile across development, others only show enrichment or depletion at specific developmental stages. Profiling of Acj6 (cholinergic enriched) and Ets65A (cholinergic depleted) binding sites *in vivo* reveals that they both directly bind the *ChAT* locus, in addition to a wide spectrum of other key neuronal differentiation genes. We also show that cholinergic enriched transcription factors are expressed in mostly non-overlapping populations in the adult brain, implying the absence of combinatorial regulation of neurotransmitter fate in this context. Furthermore, our data underlines that, similar to *C. elegans*, there are no simple transcription factor codes for neurotransmitter type specification.

## Introduction

The human brain is perhaps the most complex system known to mankind. It consists of approximately 85 billion neurons (Herculano-Houzel, 2016), which possess very diverse morphologies, neurotransmitter identities, electrical properties and preferences for synaptic partners. Understanding how this diversity is generated is one of the greatest challenges in biology and can only be achieved by identifying the underlying molecular mechanisms that determine these neuronal properties. Neurotransmitters allow neurons to communicate with each other, enabling organisms to sense, interpret and interact with their environment. Fast-acting neurotransmitters include acetylcholine and glutamate, which are, in general, excitatory, and GABA, which is inhibitory (Van Der Kloot and Robbins, 1959). The function of individual neurons depends on the specific types of neurotransmitters they produce, which in turn ensures proper information flow and can also influence the formation of neural circuits (Andreae and Burrone, 2018). Therefore the proper specification of neurotransmitter fate is fundamental for nervous system development.

Model organism studies in *C. elegans*, mice and *Drosophila* have provided a wealth of information about factors and mechanisms involved in neurotransmitter specification. Comprehensive neurotransmitter maps (Hobert, 2016) and the description of terminal selector genes in *C. elegans* (Hobert, 2008) have provided important contributions to the field. These terminal selectors are transcription factors (or a transcription factor complex) that regulate the expression of a battery of terminal differentiation genes in the last phase of neuronal differentiation, and maintain the expression of these genes during the lifetime of a neuron (Hobert, 2008). For example, the *C. elegans* transcription factors *ttx-3* and *unc-86* act as terminal selectors in distinct cholinergic and serotonergic neuron populations, respectively (Zhang et al., 2014).

Cellular context is important for the action of these specifying factors, as misexpression of terminal selectors in other neuronal subtypes is often not sufficient to reprogram their fate (Duggan et al., 1998; Wenick and Hobert, 2004). The presence of co-factors, and likely the chromatin state, can also influence this plasticity (Altun-Gultekin et al., 2001; Patel and Hobert, 2017). Related to this, there appears to be little evidence for master regulators of cholinergic, GABAergic or glutamatergic fate (Konstantinides et al., 2018; Lacin et al., 2019; Serrano-Saiz et al., 2013). Rather, individual lineages, or subpopulations, utilise different transcription factors (or combinations of transcription factors) to specify the fast-acting neurotransmitter that they will utilise. Developmental context also plays a role in the mechanisms governing neurotransmitter specification. In *Drosophila*, early born embryonic neurons in a given lineage can use different neurotransmitters (Landgraf et al., 1997; Schmid et al., 1999). However, strikingly, each post-embryonic lineage only uses one neurotransmitter (Lacin et al., 2019), implying that specification occurs at the stem cell level during larval stages.

Neurotransmitter specification studies across different organisms have highlighted conserved mechanisms. A prominent example is the binding of the transcription factors AST-1 (*C. elegans*) and Etv1 (vertebrates) to a phylogenetically conserved DNA motif to specify dopaminergic fate (Flames and Hobert, 2009). Furthermore, orthologues *acj6* (*Drosophila*), *unc-86* (*C. elegans*) and *Brn3A*/*POU4F1* (vertebrates) all have roles in cholinergic specification (Lee and Salvaterra, 2002; Serrano-Saiz et al., 2018; Zhang et al., 2014), while *PITX2* (vertebrates) and *unc-30* (*C. elegans*) both control GABAergic differentiation (Jin et al., 1994; Waite et al., 2011; Westmoreland et al., 2001).

In order to identify novel candidate genes, and investigate the dynamics of neurotransmitter specific transcription factors throughout development, we have performed cell specific profiling of RNA polymerase II occupancy, *in vivo*, in cholinergic, GABAergic and glutamatergic neurons of *Drosophila*. We identify 86 transcription factors that show differential expression between neurotransmitter types, in at least one developmental time point. There are both uniquely enriched and uniquely depleted transcription factors, and we show that acj6 (cholinergic enriched) and Ets65A (cholinergic depleted) both directly bind the choline acetyltransferase gene (*ChAT*) required for cholinergic fate.

## Results

### Transcriptional profiling of neuronal types across development

In order to investigate which genes participate in the specification of neuronal properties, namely, neurotransmitter choice, we applied the cell specific profiling technique Targeted DamID (TaDa). Targeted DamID is based on DamID (van Steensel and Henikoff, 2000) and allows the profiling of protein-DNA interactions without the need for cell isolation, specific antibodies or fixation (Aughey et al., 2019; Southall et al., 2013). Transcriptional profiling is also possible with TaDa using the core subunit of RNA polymerase II (Pol II) (Southall et al., 2013). We have mapped the occupancy of Pol II in cholinergic, GABAergic and glutamatergic neurons, using specific GAL4 drivers that trap the expression of the genes *ChAT* (choline acetyltransferase), *Gad1* (Glutamic acid decarboxylase 1) and *VGlut* (vesicular Glutamate transporter) (Diao et al., 2015). During *Drosophila* development, there are two neurogenic periods, the first to produce the larval nervous system, and the second to produce the adult nervous system. Therefore, to cover both developing stages and adult neurons, we profiled embryonic neurons, larval postembryonic neurons and adult neurons (see Figure 1B). Windows of 20 hr (embryo samples), and 24 hr (third instar larvae and adult samples) were used for TaDa profiling and 3 replicates were performed for each experiment. The number of genes bound by Pol II ranged from 1170 to 1612 (see Table S1). To investigate the global differences in Pol II occupancy between neuronal types and developmental stages, we generated a correlation matrix (Figure 2A). We found that the greatest variability is between developmental stages, rather than between cell types, with the adult brain data being more distinct from the embryonic and larval stages. When focusing on transcription factor genes, a similar pattern is evident (Figure 2B). For each developmental stage, we identified uniquely enriched genes (i.e. genes enriched in one neurotransmitter compared to the other two neurotransmitter types) (Table S2). Encouragingly, a strong enrichment of Pol II occupancy is evident at *ChAT, Gad1* and *VGlut*, the genes encoding the key enzymes involved in the biosynthesis of these neurotransmitters (Figure S1). Transcription factors and non-coding RNAs make up a large proportion of all the enriched genes, at each developmental stage (Figure 2C). In the adult, almost a quarter (23/97) of the enriched genes are transcription factors. Other enriched genes include the immunoglobulin domain containing *beaten path* (*beat*) and *Down syndrome cell adhesion molecule* (*Dscam*) genes, which play roles in axon guidance and dendrite self-avoidance (Pipes et al., 2001; Soba et al., 2007). Glutamatergic genes include *twit* and *Dad*, both of which are known to regulate synaptic homeostasis at the neuromuscular junction (Goold and Davis, 2007; Kim and Marques, 2012). Interestingly, there is an enriched expression of MAP kinase inhibitors in cholinergic neurons (Figure 2C). Also, glutamatergic neurons express higher levels of the monoamine neurotransmitter related genes *Vmat, DAT* and *Tdc2*, while GABAergic neurons are enriched for serotonergic and dopaminergic receptors, relative to the other two fast-acting neurotransmitter types (Figure 2C). Very few genes show enrichment across all developmental stages: five for cholinergic (*ChAT, ChT, acj6, Mef2* and *sosie*), five for GABAergic (*Gad1, Dbx, vg, CG13739* and *CG14989*) and two for glutamatergic (*VGlut* and *oc*) (Table S2). There is consistent enrichment of the GAL4-trapped genes (*ChAT, Gad1* and *VGlut*) (Figure S1) that provide type specific expression for the TaDa experiments.

**Figure 1.**
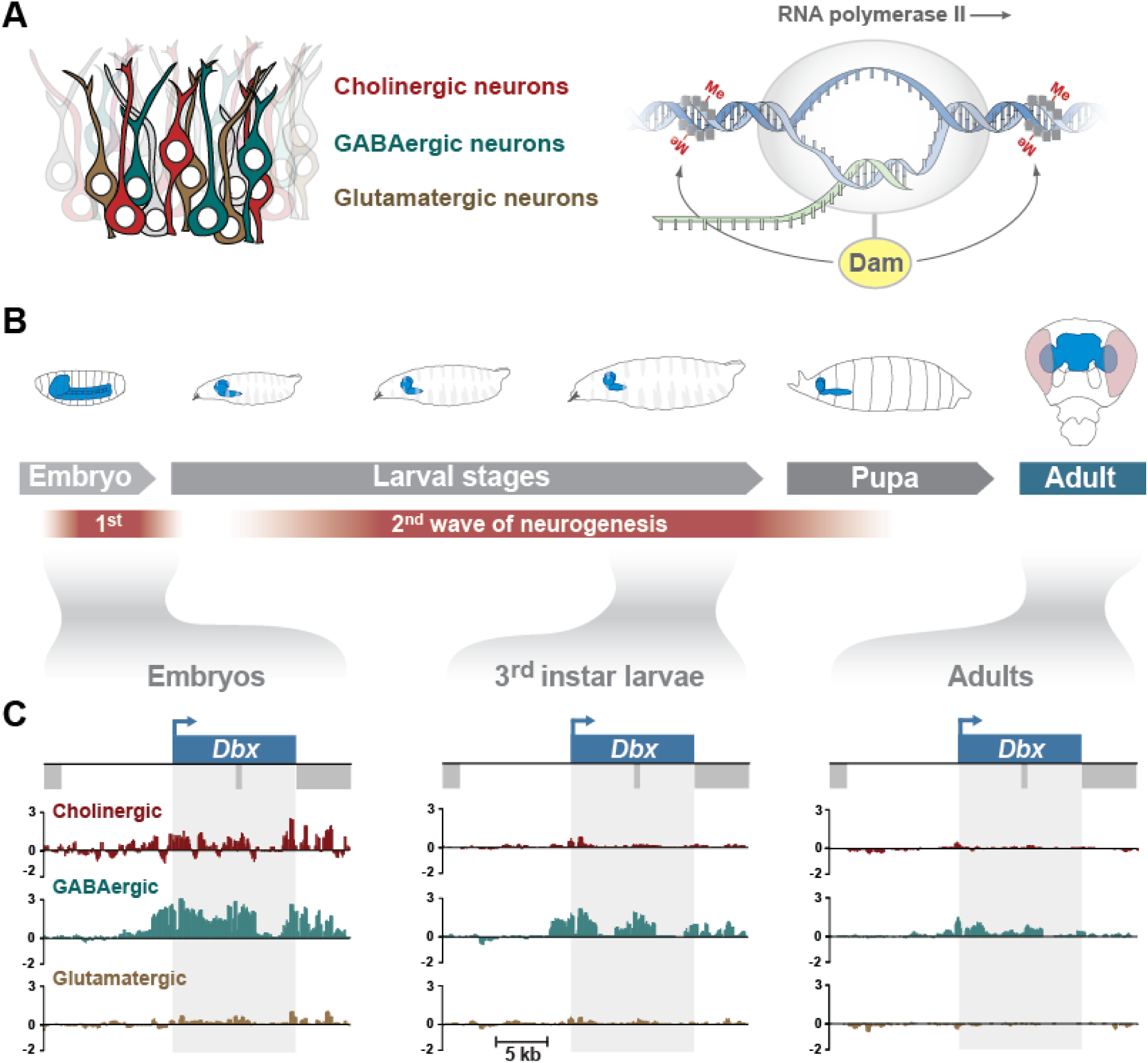
Cell specific profiling of RNA Pol II occupancy in different neuronal subtypes throughout Drosophila development. A) Profiling of RNA Pol II occupnacy in cholinergic, GABAergic and glutamatergic neurons using TaDa. B) Profling windows cover embryonic nervous system development (5 – 29 hr AEL), 3rd instar larval nervous system development (24 hr window before pupation) and the adult brain (heads from ∼ 3-4 day old adults after a 24 hr expression window). Temporal restriction of Dam-Pol II expression was controlled using a temperature sensitive GAL80. C) An example of a transcription factor gene (Dbx) that is uniquely transcribed in GABergic neurons. Y-axis represent log2 ratios of Dam-Pol II over Dam-only.

**Figure 2.**
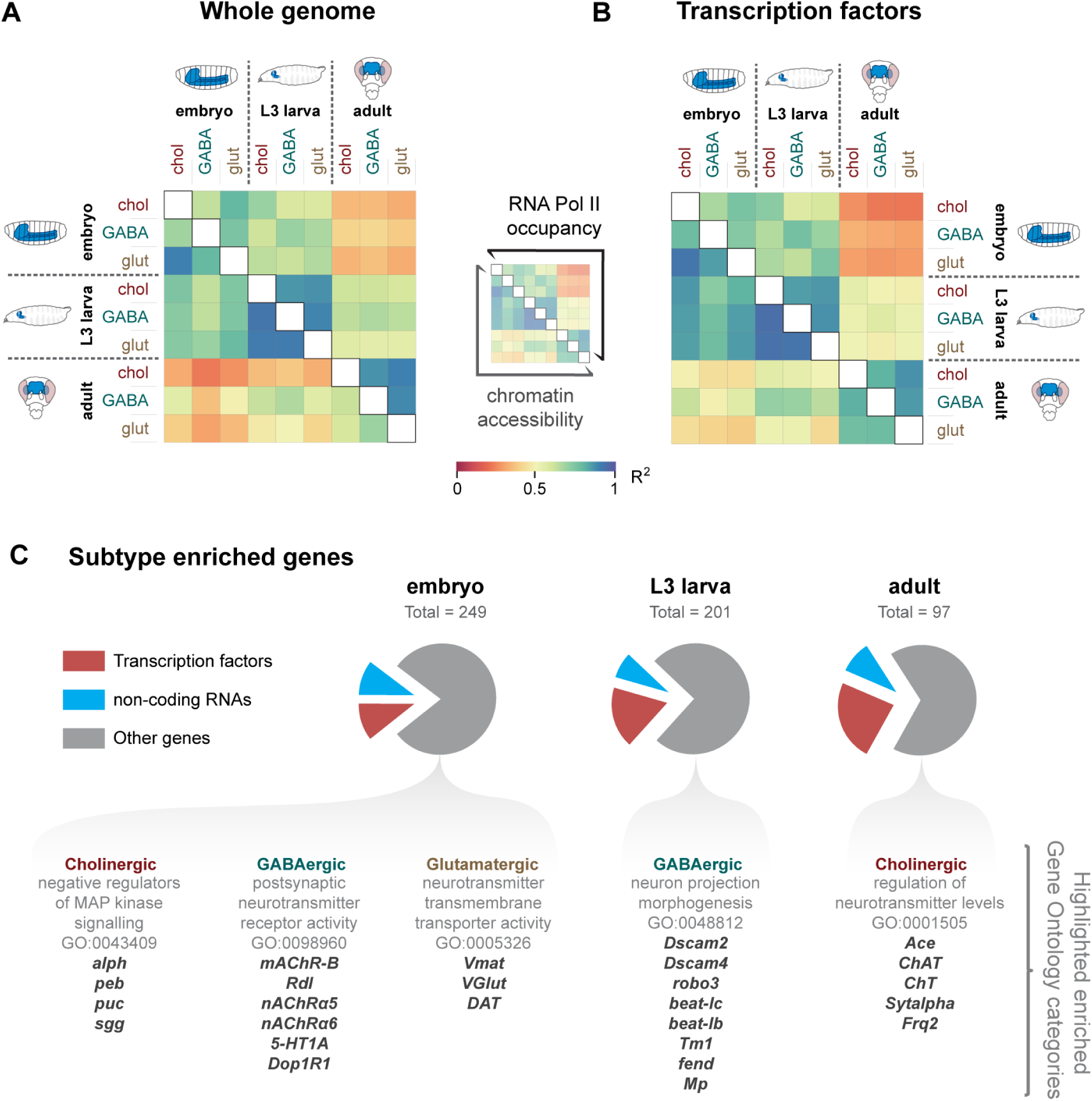
Transcription factors and non-coding RNAs are enriched in specific neurotransmitter types. A) Correlation matrix for RNA Pol II signal (log2 over Dam-only for all genes) and chromatin accessibility (CATaDa) for all gene loci (extended 5 kb upstream and 2 kb downstream). (B) Correlation matrix for RNA Pol II and chromatin accessibility at annotated transcription factor genes. (C) Proportion of transcription factors and non-coding RNAs at each developmental stage. Examples of enriched GO term categories for the remaining genes are also included.

CATaDa, an adaption of TaDa, allows profiling of chromatin accessibility without the need for cell isolation using an untethered Dam protein (i.e. the control experiment in TaDa) (Figure S2A) (Aughey et al., 2018). CATaDa reveals that, similar to RNA Pol II, global chromatin accessibility does not vary greatly between cell types (Figure 2A and 2B) but shows more differences between developmental stages. Chromatin accessibility states of embryonic and larval neurons are more similar to each other than to those of adult neurons (Figure 2A and 2B). When examining regions of the genome that display robust changes in chromatin accessibility (peaks that show >10 RPM differences across 3 consecutively methylated regions) during embryo development, only 37 GATC fragments (13 individual peaks) are identified, with 62% mapping to the loci of the three neurotransmitter synthesis genes (*ChAT, Gad1* and *VGlut*) (Figure S2B and C). This shows that across the population of neurons for each neurotransmitter type, major changes in accessibility are limited to genes involved in the respective neurotransmitter synthesis, with none of open regions directly corresponding to transcription factor loci. Differential accessibility is also present at sites outside of the gene and promoter for *Gad1* and *VGlut* (yellow arrows in Figure S2C). Weaker differences in accessibility are also observed at some of the differentially expressed transcription factor loci (Figure S2D).

### Identification of transcription factors uniquely enriched, or uniquely depleted in neurotransmitter types

Transcription factors play the major role in neurotransmitter specification and we have identified many with enriched Pol II occupancy in specific neurotransmitter types (Figure 2A). Uniquely enriched transcription factors are candidates for activators of neurotransmitter identity and conversely, if there is depletion (or absence) of a transcription factor from only one type, they are candidates for repressors of neurotransmitter identity. For example, a hypothetical transcription factor that represses GABAergic fate would be present in both cholinergic and glutamatergic neurons but absent from GABAergic neurons.

To investigate the expression pattern dynamics of both uniquely enriched and uniquely depleted transcription factors, we examined how their expression patterns transitioned across the stages of development (Figure 3). We observe a great deal of flux between transcription factor expression in cell types and developmental stages., Many genes are enriched in one or two of the developmental stages. For example, *kn, peb, rib*, and *ss* are cholinergic enriched in embryo and larva, but not in adults. *Dll* is an unusual case, as it is cholinergic enriched in larvae, however switches to being GABAergic enriched in adults (Figure 3 and Figure S3).

**Figure 3.**
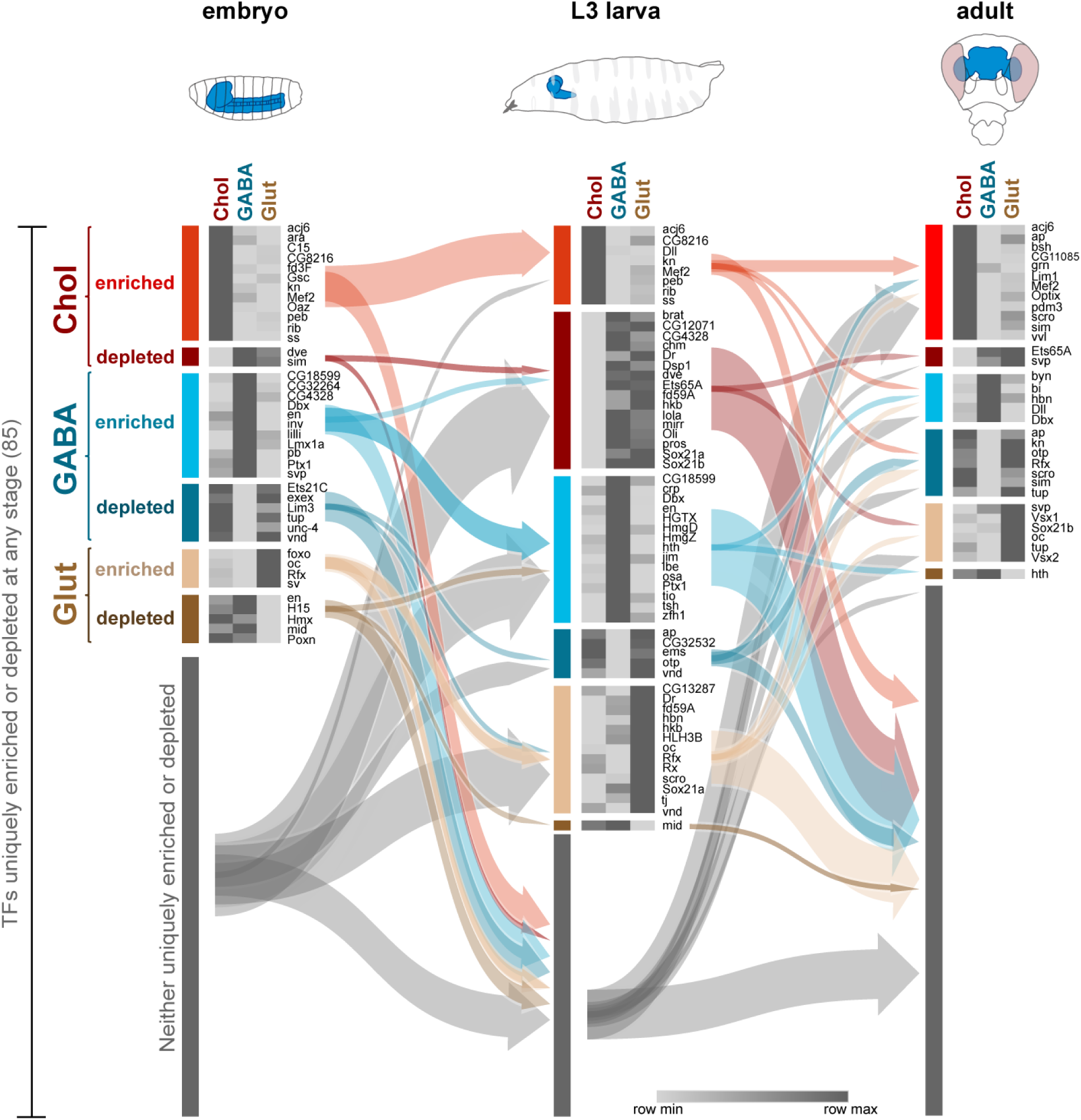
Transitions of uniquely enriched or depleted transcription factors during neural development. Transcription factors uniquely enriched or depleted in cholinergic, GABAergic and glutamatergic neurons. A total of 86 transcription factors are identified across all stages. Note that the arrows point to the group and not individual genes.

Exceptions to this are *acj6* (cholinergic – see Figure 4 and S4), *Dbx* (GABAergic – see Figure 1) and *oc* (glutamatergic), which are enriched in their respective neurotransmitter type throughout all stages. In support of our data, Acj6 is known to promote cholinergic fate in the peripheral nervous system (Lee and Salvaterra, 2002) and Dbx is important for the proper differentiation of a subset of GABAergic interneurons (Lacin et al., 2009). We checked the expression pattern of *acj6* in adult brains, and as predicted by the transcriptomic data (Figure 4B), we only found expression of acj6 in cholinergic neurons (Figure 4C). We observed the same in larval brains, with the exception of some coexpression between glutamatergic neurons and acj6 (Figure S4E). This agrees with the low level signal observed in RNA Pol II occupancy plots for *acj6* gene in third instar larva glutamatergic neurons (Figure S4A).

**Figure 4.**
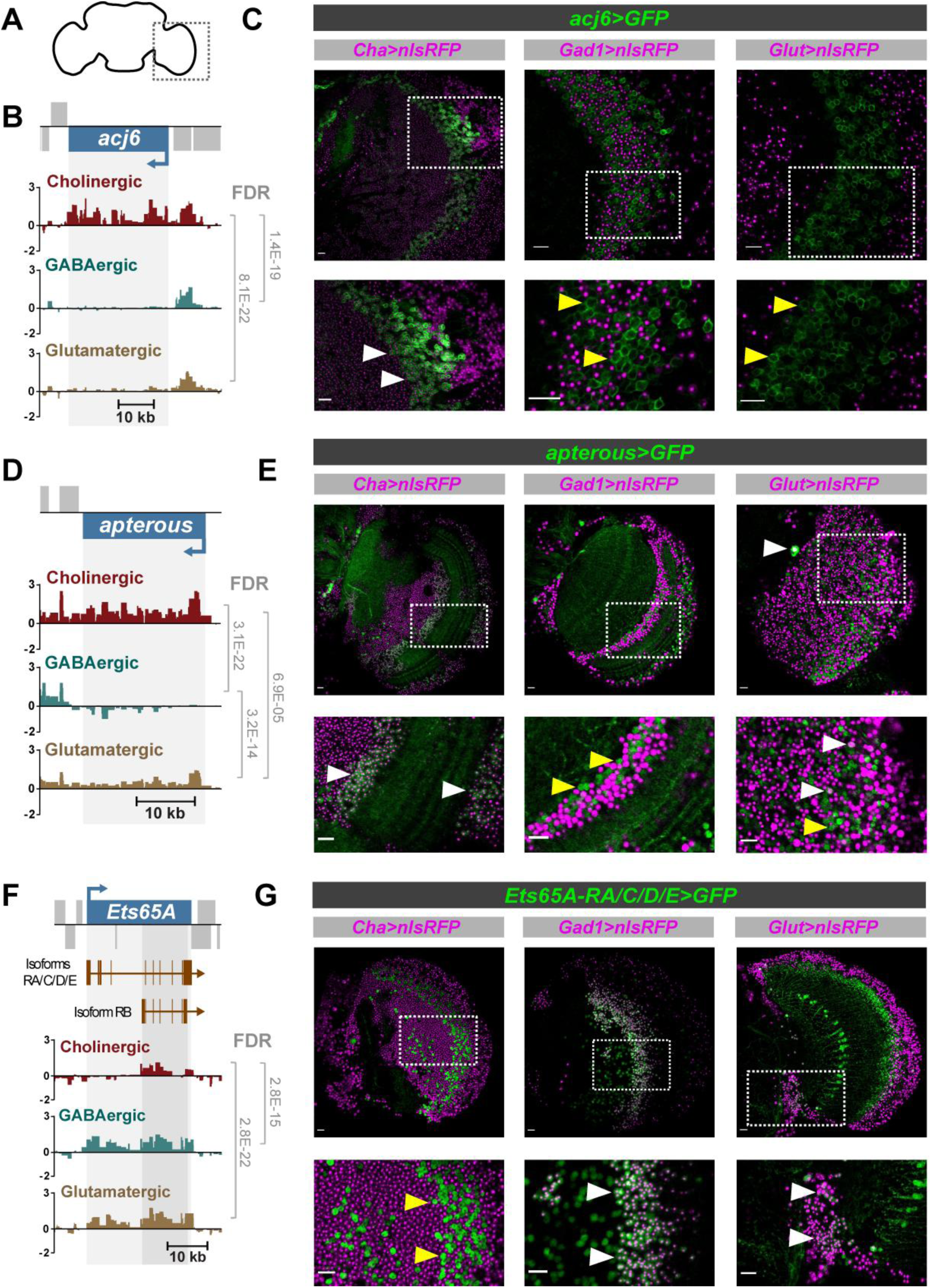
Expression of acj6, apterous and Ets65A-RA/C/D/E in the adult brain. (A) Schematic of adult brain to show region of interest. (B) Pol II occupancy at *acj6* in the adult brain. Y-axis represent log2 ratios of Dam-Pol II over Dam-only. FDR (False Discovery Rate) values are shown for significant differences (<0.01). (C) Expression pattern of *acj6*. White arrows show examples of colocalisation and yellow arrows absence of colocalisation. (D) Pol II occupancy at *apterous*. (E) Expression pattern of *apterous* in the adult brain. (F) Pol II occupancy at *Ets65A* in the adult brain. (G) Expression pattern of *Ets65A-RA/C/D/E*.

Candidate repressors of neurotransmitter fate (uniquely depleted transcription factors) also demonstrate dynamic changes in expression pattern across development (Figure 3). Prominent examples are *apterous* (absent in GABAergic), and the longer transcript isoforms of *Ets65A* (absent in cholinergic) (Figure 4D and 4F). We used genetic reporters to examine the expression pattern of *apterous* and *Ets65A-RA/C/D/E* in adult brains (Figures 4E and 4G). In agreement with our data, the GABAergic reporter is absent in *apterous* positive cells, and the cholinergic reporter is absent in *Ets65A-RA/C/D/E* positive cells. We also observed an absence of *apterous* in larval GABAergic neurons (Figure S5C and E), as predicted by the transcriptomic data (Figure S5A). As for the longer transcripts of *Ets65A-RA/C/D/E* in larval neurons, we did identify their presence in a small number of cholinergic neurons (Figure S6C and E), which could reflect the very low signal in the RNA Pol II occupancy plot (within the unique region of the long transcripts) (Figure S6A).

We have identified transcription factors with potentially novel roles in regulating neurotransmitter identity. Therefore, we investigated candidate activators and candidate repressors for their potential to elicit pan-neural reprogramming of neurotransmitter identity. Pan-neural expression and RNAi knockdown of candidate activator transcription factors (*Dbx, en, collier* and *CG4328*) and candidate repressor transcription factors (*ap, CG4328, Ets65A-RA* and *otp*) during embryonic development, and larval stages did not result in any obvious changes in neurotransmitter expression patterns (Figure S7).

Focusing on candidate transcription factors demonstrating binary differences (clear on and off), we performed literature searches to examine whether they have been previously shown, or implicated in regulating neurotransmitter identity (Figure 5). This included *C. elegans* and mouse orthologues, as much of the work in this field has utilised these model organisms. For example, the orthologues of cholinergic enriched *acj6* (*unc-86*), GABAergic enriched *Ptx1* (*PITX1* and *unc-30*) and glutamatergic enriched *oc* (*OTX1/2* and *ttx-1*) have all shown to have a role in promoting cholinergic, GABAergic and glutamatergic fate, respectively. However, there are many that have not been investigated in this context (38%), or that are only supported by indirect evidence (38%). These include *Dll* (*DLX, ceh-43*), *sox21a* (*SOX21, sox-3*), *hbn* (*ARX, alr1, unc-4*) and *otp* (*OTP, npax-1*). Given the strong conservation of neurotransmitter specification mechanisms, many of these newly highlighted factors provide promising research avenues for expanding our knowledge in this field.

**Figure 5.**
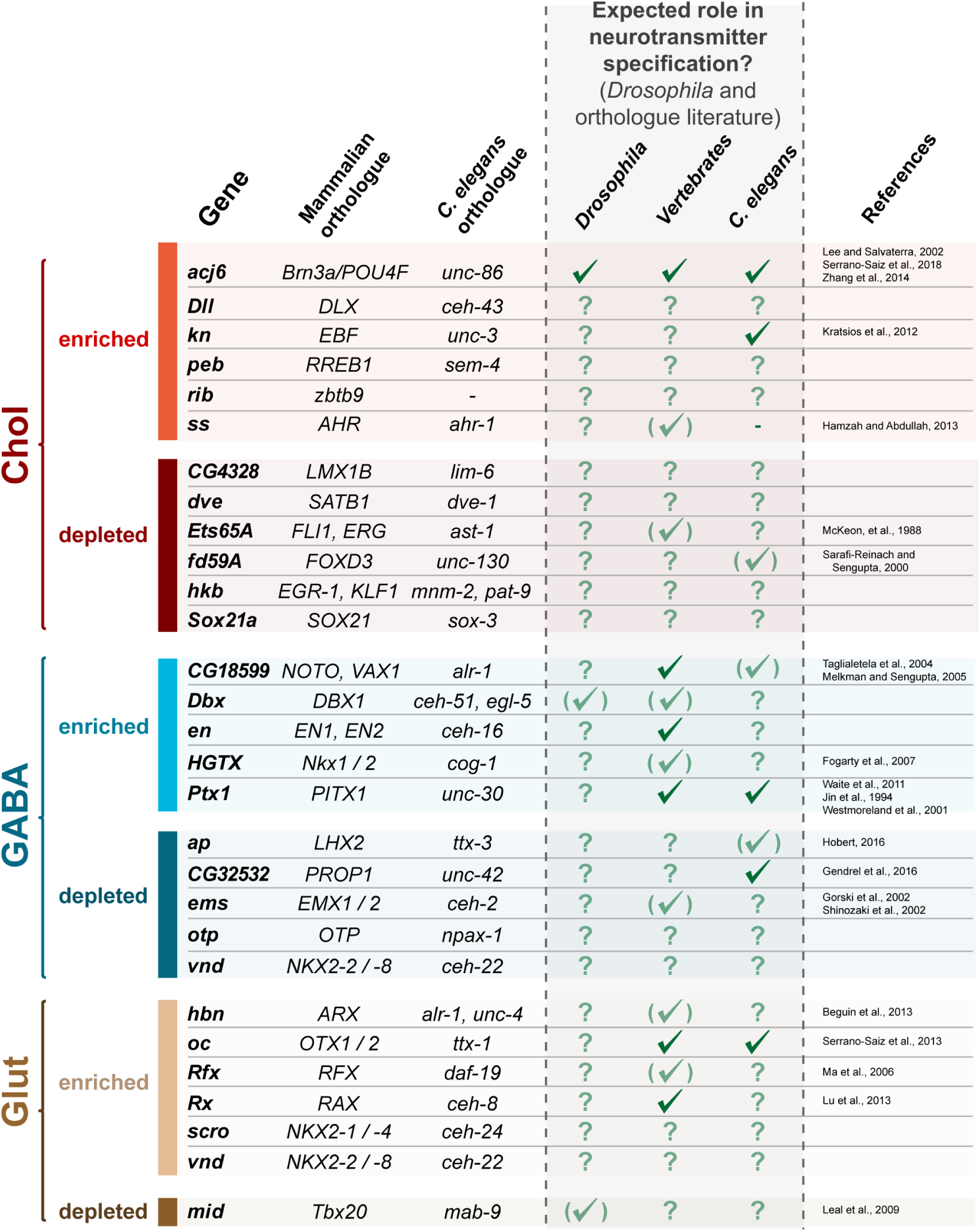
Evidence for predicted roles of identified transcription factors. Strongly enriched or depleted transcription factors identified in developing larval brains. Uniquely enriched factors are predicted to be candidates that promote the respective neurotransmitter fate, whilst uniquely depleted factors are predicted to repress the neurotransmitter fate. A full tick indicates direct evidence that the transcription factor directly promotes or represses the neurotransmitter fate, whilst indirect, supporting evidence is indicated by faded tick in brackets. A question mark signifies that nothing is currently known, regarding neurotransmitter specification.

While non-coding ribosomal RNAs and tRNAs are transcribed by RNA polymerase I and III, micro RNAs (miRNAs) and long non-coding RNAs (lncRNAs) are primarily transcribed by Pol II. Our Dam-Pol II data identifies a set of differentially bound miRNAs and lncRNAs, between the neurotransmitter types (Figure S8A). These include non-characterised lncRNAs and GABAergic enriched *iab8*, which is located in the Hox cluster and plays a role in the repression of *abd-A* (Gummalla et al., 2012). A small number of miRNAs were also identified, most notably, *mir-87* (cholinergic), *mir-184* (GABAergic) and *mir-190* (glutamatergic), which are enriched during the developing states but not in the adult. Although annotated separately, *mir-184* is embedded in *CR44206* (Figure S8B).

### Acj6 and Ets65A-PA directly bind to *ChAT* and other key neuronal differentiation genes

Acj6 is enriched in cholinergic neurons (Figure 3) and is known to promote cholinergic fate (Lee and Salvaterra, 2002). Acj6 can bind to specific sites upstream of ChAT *in vitro* (Lee and Salvaterra, 2002), however, the extent of Acj6 binding at the *ChAT* locus *in vivo*, and genome wide, is not known. In order to only profile the cells that endogenously express *acj6*, and therefore gain a more accurate readout of native Acj6 binding, we used an *acj6* GAL4 line (Lai et al., 2008) to drive the expression of the *Dam-acj6* transgene. Furthermore, we generated an *Ets65A-RA/C/D/E* MiMIC GAL4 trap line to investigate the *in vivo* binding of Ets65A-PA, with an interest to see whether, as a candidate cholinergic repressor, it could directly bind the *ChAT* locus. In the adult brain, both factors directly bind the ChAT locus (Figure 6A). Acj6 binds at the upstream region studied by (Lee and Salvaterra, 2002), as well as strongly within intronic regions of *ChAT*. Ets65A-PA also binds at the same intronic region, however, its binding at the upstream region and transcriptional start site of *ChAT* is far more pronounced (Figure 6A), which may reflect a different mode of regulation.

**Figure 6.**
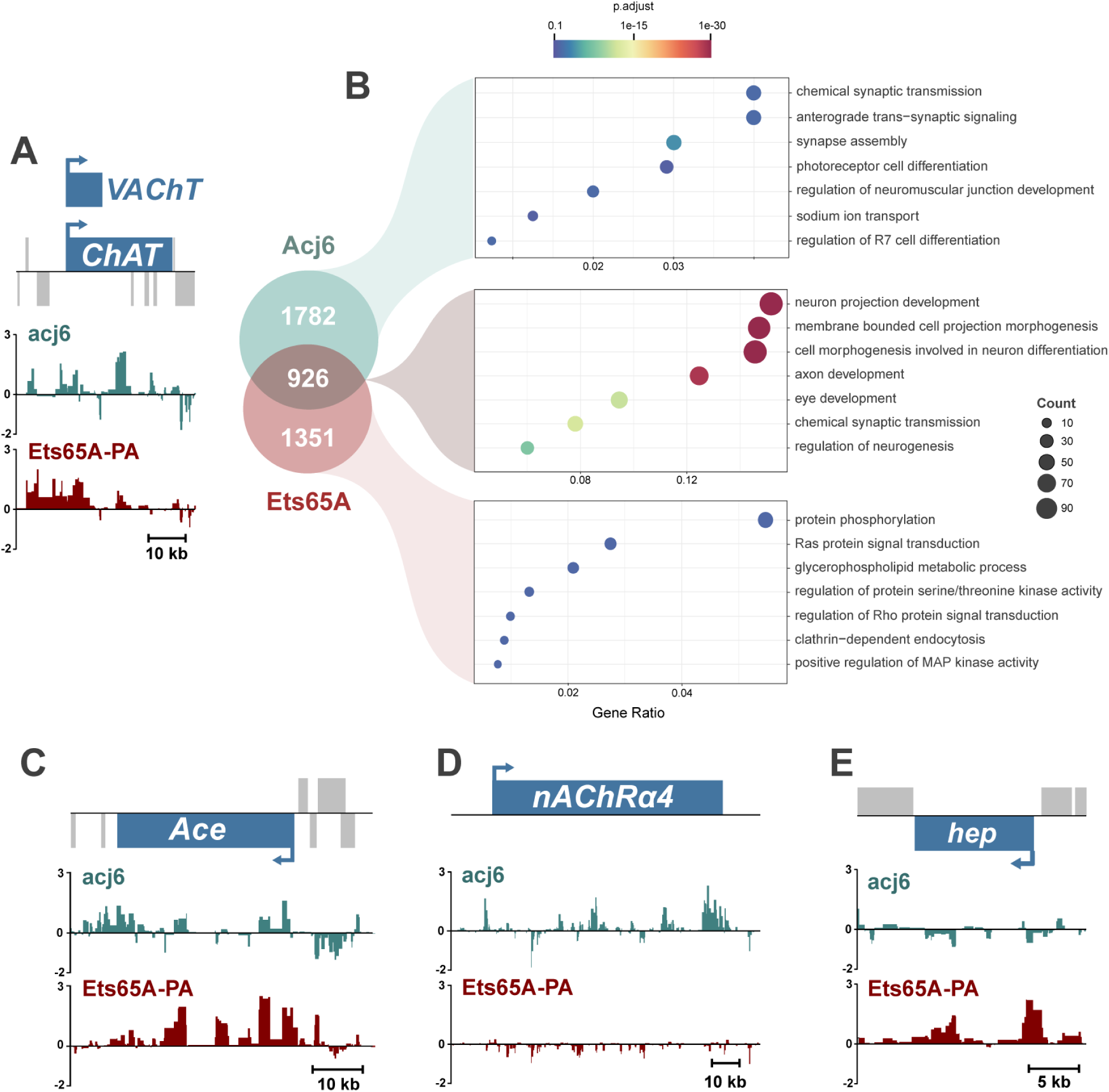
Acj6 and Ets65A-PA co-bind both *ChAT* and a whole suite of genes involved in neuronal differentiation. (A) Acj6 and Ets65-PA binding at *ChAT* (Y-axis represent log2 ratios of Dam-Pol II over Dam-only). (B) Enriched GO term categories for Acj6 and Ets65A-PA bound genes. (C)-(E) Binding at *Ace, nAChRα4* and *hep*.

Acj6 and Ets65A-PA bind 2708 and 2277 genes, respectively, using a stringent false discovery rate (FDR) (FDR < 0.0001) (Table S4). They co-bind 926 genes, which are highly enriched for nervous system genes, including genes involved in axon development [GO:0061564] and chemical synaptic transmission [GO:0007268] (Figure 6B). While both factors bind the cholinergic signalling regulator gene *Acetylcholine esterase* (*Ace*) gene (Figure 6C), Acj6 uniquely binds *nicotinic Acetylcholine Receptor α4* (*nAChRα4*) (Figure 6D), and Ets65A-RA binds multiple genes involved in MAP kinase signalling (e.g. *hep, lic, Dsor* and *slpr*) (Figure 6E). Therefore, these factors have the potential to regulate not just a single neuronal property, but also a multitude of other genes that govern a wide spectrum of neuronal processes, such as their receptivity to extrinsic signals and synapse formation.

### Enriched transcription factors are expressed in mostly non-overlapping populations

There are multiple transcription factors that show enriched expression in adult cholinergic neurons (Fig. 3). To investigate whether these factors are co-expressed within the cholinergic population, we mined single cell RNA-seq (scRNA-seq) data from adult brains (Davie et al., 2018). We find that the relative expression of the enriched factors, across the different neurotransmitter types, shows the same pattern, with enriched cholinergic factors also showing enrichment in the scRNAseq data (Figure 7A). Due to the nature of scRNAseq data, we could then determine if the cholinergic cells expressing an enriched transcription factor also express other transcription factors identified as being enriched (Figure 7B). Interestingly, there is relatively little overlap, demonstrating that these factors are expressed in distinct subpopulations of the cholinergic neurons in the adult brain.

**Figure 7.**
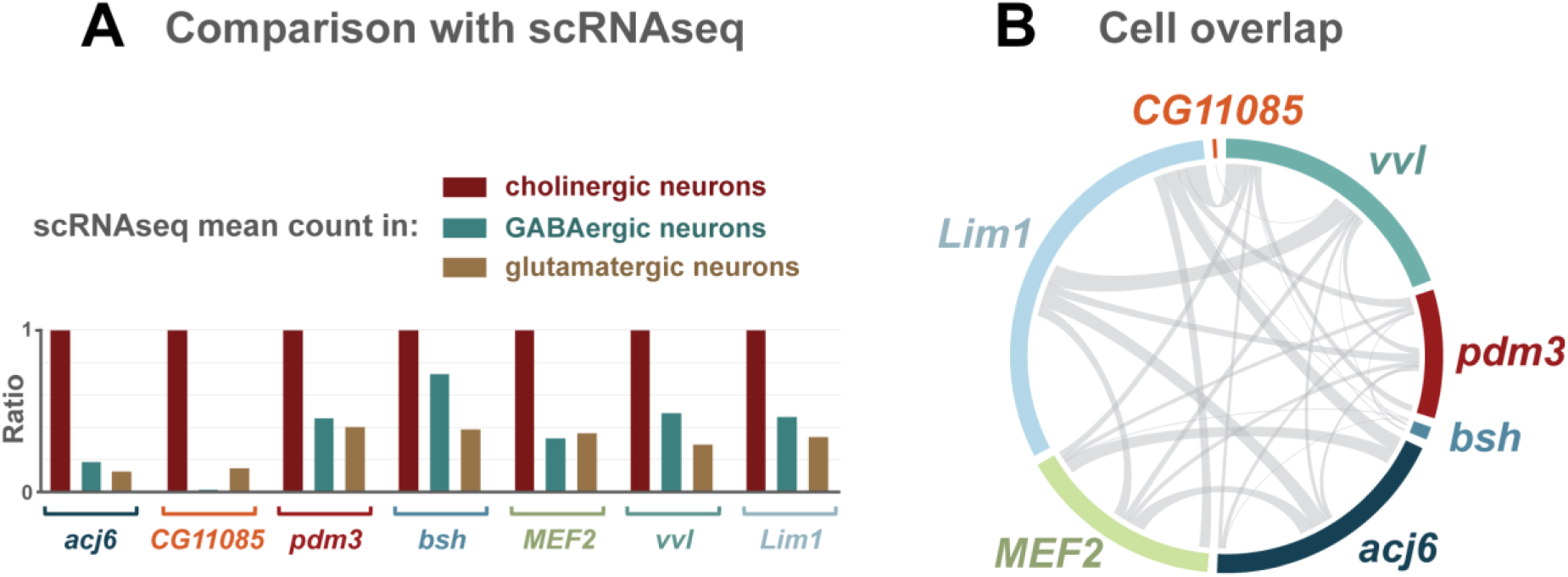
Enriched transcription factors are expressed in mostly non-overlapping populations of adult cholinergic neurons. A) Transcription factors identified as enriched in cholinergic neurons by Targeted DamID are also enriched in scRNAseq data (adult brain). Mean counts are ratio normalised to the average count value in cholinergic neurons. B) Circos plot displaying the overlap in cells expressing cholinergic enriched transcription factors.

## Discussion

Neurotransmitter identity is a key property of a neuron that needs to be tightly regulated in order to generate a properly functioning nervous system. Here we have investigated the dynamics and extent of transcription factor specificity in fast-acting neurotransmitter neuronal types in *Drosophila*. We profiled the transcription state of cholinergic, GABAergic and glutamatergic neurons in the developing and adult brain of *Drosophila*. We observe enriched Pol II occupancy at the relevant neurotransmitter synthesis genes (Figure S1) and other genes associated with the activity of the specific types (Table S2). The monoamine neurotransmitter related genes *Vmat, DAT* and *Tdc2* are enriched in glutamatergic neurons (Figure 2C), which is not unprecedented, as monoamine populations can also be glutamatergic (Aguilar et al., 2017; Trudeau and El Mestikawy, 2018). Cholinergic, GABAergic, serotonergic and dopaminergic receptors are enriched in embryonic GABAergic neurons relative to the other two fast-acting neurotransmitter types (Figure 2C), which correlates with GABAergic interneurons acting as integrative components of neural circuits. The enrichment of MAP kinase pathway regulators in cholinergic neurons is intriguing, suggesting that this signalling pathway may have a specific role in these neurons. This is supported by a recent study showing that MAP kinase signalling acts downstream of Gq-Rho signalling in *C. elegans* cholinergic neurons to control neuron activity and locomotion (Coleman et al., 2018).

Importantly, we have uncovered and highlighted transcription factors and non-coding RNAs differentially expressed between these types. Some of these are expected based on previous studies in *Drosophila*, including *acj6* (cholinergic) (Lee and Salvaterra, 2002) and *Dbx* (GABAergic) (Lacin et al., 2009). Also, studies in other model organisms fit with our findings, for example, cholinergic enriched *knot*, whose orthologue, *UNC-3* (*C. elegans*), is a terminal selector for cholinergic motor neuron differentiation (Kratsios et al., 2011). In addition, *RFX*, the vertebrate orthologue of *Rfx*, which we identified as glutamatergic enriched, can increase the expression of the neuronal glutamate transporter type 3 (Ma et al., 2006). However, we have identified many differentially expressed transcription factors that have not had their role studied with respect to neurotransmitter specification, or cases where there is supportive, but not direct, evidence for a role in neurotransmitter specification. For instance, vertebrate neuronal precursors expressing *Nkx2*.*1 (HGTX* orthologue) predominantly generate GABAergic interneurons (Fogarty et al., 2007), and a polyalanine expansion in *ARX* (*hbn* orthologue) causes remodelling and increased activity of glutamatergic neurons in vertebrates (Beguin et al., 2013). *Acj6* is expressed in a subset of cholinergic neurons (Lee and Salvaterra, 2002) and *Dbx* in a subset of GABAergic neurons (Lacin et al., 2009). To the best of our knowledge, none of the enriched transcription factors we identified are expressed in all of the neurons of a particular neurotransmitter type. This highlights that, similar to *C. elegans* (Hobert, 2016), there are no simple transcription factor codes for neurotransmitter type specification in *Drosophila*.

Uniquely enriched factors are candidates for promoting a neurotransmitter fate, and we tested a number of them for their ability to reprogram neurons on a global scale in embryos (Figure S7). No obvious changes were observed, however, this is not particularly surprising considering the importance of cellular context for the reprogramming of neuronal properties (Duggan et al., 1998; Wenick and Hobert, 2004). Successful reprograming may require intervention at a specific time point (e.g. at the progenitor stage), the co-expression of appropriate co-factors, and/or to exclusively target a neuronal subpopulation within each neurotransmitter type. Future work could investigate these factors in specific and relevant lineages, to shed light on important contextual information.

The majority of transcription factors identified as directly regulating neurotransmitter fate act in a positive manner, whereas only a handful of studies describe the role of repressors. Incoherent feedforward loops exist in *C. elegans*, where terminal selectors activate repressors, which feedback onto effector genes (for review, see (Hobert, 2016)). In vertebrates, both *Neurogenin 2* and *Tlx3* are required for the specification of certain glutamatergic populations but also act to repress GABAergic fate (Cheng et al., 2004; Schuurmans et al., 2004). Whether this is direct repression of *Glutamic acid decarboxylase* (*Gad*) genes (required for the synthesis of GABA), or indirectly, through another transcription factor, is unclear. We have identified several transcription factors that are expressed in two neurotransmitter types, but absent from the other. These include *apterous* (*ap*), *Ets65A* (long transcripts) and *orthopedia* (*otp*), which we hypothesise to be candidate repressors, given their absence from cells with a specific neurotransmitter identity. Our profiling of Ets65A-PA binding *in vivo*, reveals that it directly binds *ChAT* (Figure 6A), and therefore has the potential to directly regulate cholinergic fate. Similar to the candidate activators, ectopic expression of these candidates did not show any obvious repression of the respective neurotransmitter genes (Figure S7), however, again, this might be because they can only act as a repressor in specific contexts (e.g. when a co-repressor is present), or that they regulate genes associated with specific types but do not directly regulate neurotransmitter identity.

The development of single cell RNA-seq (scRNA-seq) technology has led to the profiling of several *Drosophila* tissues, including the whole adult brain (Davie et al., 2018), the central adult brain (Croset et al., 2018) and the adult optic lobes (Konstantinides et al., 2018). Here we mined the whole adult brain data (Davie et al., 2018) to compare and investigate the cholinergic enriched factors that we identified in adult brains. The enrichment of these transcription factors (compared to GABAergic and glutamatergic neurons) is also observed in the scRNAseq data (Figure 7A). Furthermore, we discovered that the cholinergic cells that these factors are expressed in are almost non-overlapping (Figure 7B). This is an intriguing finding, as it suggests that these factors, if they are indeed acting to promote/maintain cholinergic fate, they are not acting together in this context. This scenario maybe different during development, where specification is occurring, and it will be interesting to test this when high coverage scRNAseq data is available for the 3^rd^ instar larval brain. We observed more differentially expressed transcription factors in the L3 larval stage (58) compared to the embryo (40) or adults (33). This may reflect the existence of both the functioning larval nervous system (built during embryogenesis) and the developing adult nervous system at this stage (Figure 3). While both the embryo and larval data are similar on a global scale, Pol II occupancy and chromatin accessibility in the adult brain is less correlated (Figure 2). It is currently unclear whether this is due to adult VNCs being absent from the profiling experiments, or differences between immature and fully mature neurons, such as overall lower transcriptional activity in adults. We have previously shown that global chromatin accessibility distribution in adult neurons is distinct from larval neurons (Aughey et al., 2018), which may account for some of these differences.

Apart from the neurotransmitter synthesis genes, the chromatin accessibility of the different neuronal types, at a given stage, is surprisingly similar, as demonstrated in embryos (Figure S2B). The enriched accessibility is not just restricted to the gene bodies of the neurotransmitter genes, and peaks are present upstream (*Gad1*) and downstream (*VGlut*) (Figure S2C), which are likely enhancers. Accessibility at the *ChAT* gene is clearly higher in cholinergic neurons at the embryonic and adult stages, however, in third instar larvae, the difference is less pronounced (Figure S2C). This could reflect increased plasticity at this stage, possibly linked to the dramatic remodelling of larval neurons during metamorphosis (for a review, see (Yaniv and Schuldiner, 2016)). While a subset of transcription factors display obvious contrasts in Pol II occupancy, the same transcription factors have no observable, or minor, differences in accessibility (Figure S2D). This could be due to transcription factors being expressed at relatively lower levels and/or that they are only expressed in a subset of the cells, therefore the difference is less prominent.

Evidence is emerging for the roles of miRNAs in generating neuronal diversity, including the differentiation of taste receptor neurons in worms (Chang et al., 2004; Johnston and Hobert, 2005) and dopaminergic neurons in vertebrates (Kim et al., 2007). Here, we found the enriched expression of *mir-184* in GABAergic cells (Figure S8B), which is intriguing, as *mir-184* has been shown to downregulate *GABRA3* (GABA-A receptor) mRNA (possibly indirectly) in vertebrate cell lines (Luo et al., 2017), and may be a mechanism to help prevent GABAergic neurons self-inhibiting. Furthermore, *mir-87* has enriched RNA polymerase II occupancy in cholinergic neurons (Figure S8A), and when mutated causes larval locomotion defects in *Drosophila* (Picao-Osorio et al., 2017).

*Acj6* is expressed in adult cholinergic neurons (Figure 4B and C)(Lee and Salvaterra, 2002), whilst *Ets65A-PA* is expressed in non-cholinergic adult neurons (Figure 4F and G). However, despite this, they bind a large number of common target genes (Figure 6). This includes 20% (101/493) of all genes annotated for a role in *“neuron projection development”* (GO:0031175). This is quite striking, especially as this is in the adult, where there is virtually no neurogenesis or axonogenesis. However, this may reflect dendritic re-modelling processes, or a requirement of neurons to continuously express transcription factors, even after development, to maintain their fate. The *acj6* orthologues, *unc-86* and *Brn3a* are both required to maintain the fate of specific cholinergic populations (Serrano-Saiz et al., 2018), and transcriptional networks that specific Tv1/Tv4 neurons in *Drosophila* are also required to maintain them in the adult (Eade et al., 2012). Therefore, the binding of Acj6 and Ets65A-PA to developmental genes and *ChAT* in adult neurons could be required for the continued activation (and repression) of genes governing neuronal identity. MAP kinase signalling genes are enriched in cholinergic neurons (Figure 2C) and Ets65A-PA specifically binds MAP kinase signalling genes (Figure 6), making it tempting to speculate that Ets65A-PA acts to repress cholinergic specific genes such as *ChAT* and MAP kinase genes. These Acj6 and Ets65A-PA data also emphasise the diverse set of neuronal differentiation genes a single transcription factor could regulate.

The precise synthesis and utilisation of neurotransmitters ensures proper information flow and circuit function in the nervous system. The mechanisms of specification are lineage specific, predominantly through the action of transcription factors. Here we have provided further insights into the complement of different transcription factors that regulate neurotransmitter identity throughout development. Furthermore, we identified the genomic binding of a known activator, and a candidate repressor, of cholinergic fate in the adult, emphasising the broad spectrum of neural identity genes that they could be regulating outside of neurotransmitter use. Given the strong evidence for conserved mechanisms controlling neurotransmitter specification, these data will be a useful resource for not just researchers using *Drosophila* but other other model systems too. Continued work to elucidate the mechanisms, co-factors and temporal windows in which these factors are acting will be fundamental in gaining a comprehensive understanding of neurotransmitter specification.

## Materials and Methods

### *Drosophila* lines

Lines used in this study are as follows:

*w; dvGlut-GAL4 [MI04979]/CyO act-GFP*, (Bloomington #60312)

*w;; ChAT-GAL4 [MI04508] / TM3 act GFP*, (Bloomington #60317)

*w;; Gad1-GAL4 [MI09277] / TM3 actin GFP* (Diao et al., 2015)

*w[*]; Mi{Trojan-lexA:QFAD*.*2}VGlut[MI04979-TlexA:QFAD*.*2]/CyO, P{Dfd-GMR-nvYFP}2*, (Bloomington #60314)

*w[*]; Mi{Trojan-lexA:QFAD*.*0}ChAT[MI04508-TlexA:QFAD*.*0]/TM6B, Tb[1]*, (Bloomington #60319)

*w[*]; Mi{Trojan-lexA:QFAD*.*2}Gad1[MI09277-TlexA:QFAD*.*2]/TM6B, Tb[1]*, (Bloomington, #60324). (All obtained from M. Landgraf)

*UAS-LT3-NDam, tub-GAL80*^*ts*^; *UAS-LT3-NDam-RNA Pol II* (from Andrea Brand)

*Ets65A-RA/C/D/E-GAL4 [MI07721]* (this study)

*apterous-GAL4; UAS-GFP* (from F Jiménez Díaz-Benjumea)

*acj6-GAL4-UAS-mCD8-GFP/FM7c; Pin/CyO* (from DJ Luginbuhl) (Lai et al., 2008)

*elavG4;; Mi{PT-GFSTF*.*2}Gad [MI09277]/TM3 actin-GFP* (Bloomington, #59304)

*elavG4;; Mi{PT-GFSTF*.*0}ChAT [MI04508]/TM3 actin-GFP* (Bloomington, #60288)

*UAS-Dbx* (Bloomington, #56826)

*UAS-apterous* (from F Jiménez Díaz-Benjumea),

UAS-collier (from F Jiménez Díaz-Benjumea),

*UAS-engrailed [E9]* (from Andrea Brand)

*UAS-otp* (Fly ORF #F000016)

*UAS-CG4328* (FlyORF, #F0019111)

*UAS-Dbx sh RNAi attP40* (VDRC #330536)

*UAS-ap sh RNAi attP40* (VDRC #330463)

*UAS-Ets65A-RA RNAi attP2* (Bloomington #41682)

*UAS-Ets65A-RA attP2* (this study)

*yw, hs-Flp 1; +; Dr/TM6B*

*yw, hs-Flp 1; +; UAS-Ets65A-RA*

*AyGal4, UAS-mCD8-GFP/(CyO); Cha lexAQF, mCherry /TM6B*

### Generation of *Ets65A* and *acj6* Targeted DamID lines

Details and sequences of all primers used for generating constructs are shown in Supplemental Experimental Procedures. *pUAST-LT3-NDam-acj6-RF* and *pUAST-LT3-NDam-Ets65A-RA* were generated by PCR amplifying *acj6-RF* and *Ets65A-RA* from an embryonic cDNA library. The resulting PCR products were cloned into *pUAST-LT3-Dam* plasmid (Southall et al., 2013) with NotI and XhoI sites, using Gibson assembly.

acj6-RF FW:

CATCTCTGAAGAGGATCTGGCCGGCGCAGATCTGCGGCCGCTCATGACAATGTCGATGTATTCGACGACGG

acj6-RF RV:

GTCACACCACAGAAGTAAGGTTCCTTCACAAAGATCCTCTAGATCAGTATCCAAATCCCGCCGAACCG

Ets65A-RA FW:

CTGAAGAGGATCTGGCCGGCGCAGATCTGCGGCCGCTCATGTACGAGAACTCCTGTTCGTATCAGACG

Ets65A-RA RV:

ACAGAAGTAAGGTTCCTTCACAAAGATCCTCTAGATCATGCGTAGTGGGGATAGCTGCTC

### Generation of *Ets65A-RA-GAL4* line

*Ets65A-RA/C/D/E-GAL4* was generated by inserting a GAL4 trap cassette into the MI07721 MiMIC line (Bloomington #43913) using the triplet donor *in vivo* system described in (Diao et al., 2015).

### Targeted DamID (TaDa) for RNA-Pol II mapping

Crosses producing larvae with the following genotypes were allowed to lay eggs over a minimum of two days at 25°C before timed collections were performed:

*tub-GAL80*^*ts*^*/+; UAS-LT3-NDam/ ChAT-GAL4*^*MI04508*^

*tub-GAL80*^*ts*^*/+; UAS-LT3-NDam-RNA Pol II/ ChAT-GAL4*^*MI04508*^

*tub-GAL80*^*ts*^*/+; UAS-LT3-NDam/ Gad1-GAL4* ^*MI09277*^

*tub-GAL80*^*ts*^*/+; UAS-LT3-NDam-RNA Pol II/ Gad1-GAL4* ^*MI09277*^

*tub-GAL80*^*ts*^*/ dvGlut-GAL4* ^*MI04979*^; *UAS-LT3-NDam/ +*

*tub-GAL80*^*ts*^*/ dvGlut-GAL4* ^*MI04979*^; *UAS-LT3-NDam-RNA Pol II/ +*

### First instar larvae samples

Crosses of the right genotype were allowed to lay old eggs for 2 hours at 25°C in fly cages. Wet yeast and two drops of 10% acetic acid were added to apple juice plates to promote egg laying. Then, egg laying was done for 5 hours at 25°C, apple juice plates containing those embryos were transferred to 29°C (permissive temperature) for 20 hours. After this time, first instar larvae were collected and stored in 1x PBS. Samples were flash-frozen in dry ice, and stored at −80°C till the appropriate amount of tissue was enough to start the experiment. No selection for the right genotype was done, and 20 µl worth of volume of tissue was used as a proxy to determine the appropriate amount of material for each replicate. 3 replicates were done for each experiment. With this husbandry protocol, the collected first instar larvae were around 12 hours ALH (after larvae hatching), just before the first larval neurons are being generated, then providing the transcriptome of embryonic neurogenesis.

### Third instar larvae samples

Crosses of the right genotypes were allowed to lay eggs for 6 hours at 25°C in fly food vials. These vials were then transferred to 18°C (restrictive temperature) for 7 days. They were then moved to 29°C (permissive temperature) for 24 hours. Wandering stage larvae, around 96h ALH, were selected with a GFP scope for the right genotype. Larvae were dissected in 1x PBS, leaving the anterior half of the larvae partly dissected, containing the CNS, but removing the gut and all the fat tissue. Samples were flash-frozen in dry ice, and stored at −80°C till the appropriate amount of tissue was enough to start the experiment. 100 partly dissected CNS were used for each replicate. 3 replicates were done for each experiment.

### Adult samples

Crosses of the right genotypes were allowed to lay eggs for 2 days at 18°C in fly food vials. Vials containing those eggs were kept at 18°C (restrictive temperature) till adult flies eclosed. They were then kept at 18°C for 5-10 days. After that, they were selected according for the right genotype, and transferred to 29°C (permissive temperature) for 24 hours. Then, adult flies were flash-frozen in dry ice, and stored at −80°C. Around 50 fly heads were used for each replicate. 3 replicates were done for each experiment.

When preparing the tissue to be used, larvae nor flies were sex sorted. It has been recently reported that transcriptomes from males and females, obtained with a cell specific driver combination expressed in neurons in the adult optic lobe, do not present major differences in their transcriptomes. Only a small number of genes, known sex-specific genes showed differences between sexes. (Davis et al., 2018).

Our DamID protocol was based on (Southall et al., 2013), and (Marshall et al., 2016). Briefly, DNA was extracted using Qiagen DNeasy kit, and a minimum of 3 µg of DNA was precipitated for first instar larvae, 6ug for third instar larvae, and 2.5 µg for adult samples. DNA was digested with DpnI overnight at 37°C. The next morning, 0.5 µl of DpnI was added for 1 hour extra incubation, followed by DpnI heat inactivation (20 mins 80°C). Either Advantage cDNA polymerase, or Advantage 2 cDNA polymerase mix, 50x, Clontech, were used in PCR amplification. Enzymes Sau3AI or AlwI were used to remove DamID adaptors, from sonicated DNA.

Libraries were sequenced using Illumina HiSeq single-end 50 bp sequencing. Three replicates were performed for each experiment. A minimum of 25 million reads were obtained from the first instar larvae samples, 30 million reads from the third instar larvae, and 9 million reads from the adults’ samples.

### Targeted DamID (TaDa) for identification of acj6 and Ets65A-RA binding sites

Crosses producing larvae with the following genotypes were allowed to lay eggs over a minimum of two days at 25°C:

*acj6-GAL4-UAS-GFP; tub-GAL80*^*ts*^*/+; UAS-LT3-NDam/+*

*acj6-GAL4-UAS-GFP; tub-GAL80*^*ts*^*/+; UAS-LT3-NDam-acj6-RF/+*

*tub-GAL80*^*ts*^*/+; UAS-LT3-NDam/ Ets65A-RA-GAL4*^*MI07721*^

*tub-GAL80*^*ts*^*/+; UAS-LT3-NDam-Ets65A-RA/ Ets65A-RA-GAL4*^*MI07721*^

Crosses of the right genotypes were allowed to lay eggs for 2 days at 18°C in fly food vials. Vials containing those eggs were kept at 18°C (restrictive temperature) till adult flies eclosed. They were then kept at 18°C for around 10 days. Then, they were transferred to 29°C (permissive temperature) for 24 hours, selected according for the right genotype, flash-frozen in dry ice, and stored at −80°C. A minimum of 150 fly heads were used for each replicate. 2 replicates were done for each experiment.

The DamID protocol used for these samples is the same as described above, with minor changes, 6 µg of DNA were precipitated, Bioline Polymerase was used in the PCR amplification, and only AlwI was used to remove adaptors from sonicated DNA. Libraries were sequenced using Illumina HiSeq single-end 50 bp sequencing. Two replicates were acquired for each experiment. A minimum of 10 million reads were obtained these samples.

### Targeted DamID data analysis

Sequencing data for TaDa and CATaDa were mapped back to release 6.03 of the *Drosophila* genome using a previously described pipelines (Aughey et al., 2018; Marshall and Brand, 2015). Transcribed genes (defined by Pol II occupancy) were identified using a Perl script described in (Mundorf et al., 2019) based on one developed by (Southall et al., 2013) (available at https://github.com/tonysouthall/Dam-RNA_POLII_analysis). *Drosophila* genome annotation release 6.11 was used, with 1% FDR and 0.2 log2 ratio thresholds. To compare data sets, log2 ratios were subtracted, in this case, producing 3 replicate comparison files (as 3 biological replicates were performed). These data were then analysed as described above to identify genes with significantly different Pol II occupancy. Due to the presence of negative log2 ratios in DamID experiments, these genes were filtered to check that any significantly enriched genes were also bound by Pol II in the experiment of interest (numerator data set). A gene list was generated from the transcript data using the values from the associated transcript with the most significant FDR. Correlation values (Figure 2) were visualised using Morpheus (https://software.broadinstitute.org/morpheus/). The transition plot (Figure 3) was generated in R using the transitionPlot function from the Gmisc R package (http://gforge.se/). Enrichment GO analysis was performed using the R package clusterProfiler (Yu et al., 2012)

For acj6 and Ets65A-RA TaDa, peaks were called and mapped to genes using a custom Perl program (available at https://github.com/tonysouthall/Peak_calling_DamID) In brief, a false discovery rate (FDR) was calculated for peaks (formed of two or more consecutive GATC fragments) for the individual replicates. Then each potential peak in the data was assigned a FDR. Any peaks with less than a 0.01% FDR were classified as significant. Significant peaks present in all replicates were used to form a final peak file. Any gene (genome release 6.11) within 5 kb of a peak (with no other genes in between) was identified as a potentially regulated gene.

For studying transcription factors specifically, we filtered the differentially expressed genes for known/predicted transcription factors using the FlyTF database (Pfreundt et al., 2010).

### Extracting gene specific data from scRNAseq data

Data for specific genes were extracted from the adult scRNAseq matrix file (Davie et al., 2018) using the following Perl code:

~~~
#!usr/bin/perl
#parse_scRNAseq_data
use warnings;

my $file = ‘mtx_file.mtx’; #path to the scRNAseq matrix file

print “\nEnter gene number to extract for – see gene index file\n\n”;
$genenum = <STDIN>; chomp $genenum; chomp $genenum;

open (OUTPUT, ‘> scRNAseq_data_for_gene’.”$genenum”.’.txt’);
open my $fh, ‘<‘, $file or die $!;
while(<$fh>){@col = split(/\s/,$_); if($genenum == $col[0])
    {print OUTPUT “$col[0]\t$col[1]\t$col[2]\n”;}} exit;
~~~

Cells with a transcript count (for the given gene) of less than 3 were excluded for further analysis.

### Immunostaining and imaging

Third instar larval CNS or adult brains were dissected in 1x PBS. They were fixed in 4% formaldehyde (methanol free) 0.1% Triton X-100 PBS (PBST), for 30 minutes at room temperature. Samples were then rinsed twice with 0.1% PBST, and washed four times for 1 hour with 0.1% PBST. 5% Normal Goat Serum in 0.1% PBST was used as a blocking agent for 1 hour at room temperature. Brains were then incubated overnight at 4°C with primary antibodies in 5% Normal Goat Serum in 0.1% PBST. The primary antibodies used were: anti-Chicken-GFP (Abcam #13970, 1:2000), and anti-Rabbit-DsRed (Clontech #632496, 1:500). Brains were rinsed twice with 0.1% PBST, and washed four times with 0.1% PBST for 1 hour. Secondary antibodies were diluted in 5% Normal Goat Serum in 0.1% PBST and incubated with the brains for 1 hour at room temperature. The secondary antibodies used were: anti-Chicken-Alexa 488 (Thermo Scientific #A11039, 1:500), and anti-Rabbit-Alexa 546 (Thermo Scientific #A11010, 1:500). Samples were then rinsed twice with 0.1% PBST, and washed four times for 1 hour. Brains were mounted on glass cover slides in Vectashield (Vector laboratories). All incubations and washes were performed in a rotator. After dissection of first instar larvae CNS, they were placed in a polylisine coated microscope slide, where we performed all the incubations. Both experimental CNS and wild-type CNS were placed on the same slide. For all the immunostaining experiments, a minimum of 5 brains were dissected and visualised. Images were acquired using a Zeiss LSM 510 confocal microscope and edited using Fiji/Image J.

### Accession numbers

All raw sequence files and processed files have been deposited in the National Center for Biotechnology Information Gene Expression Omnibus (accession number GSE139888).

## Acknowledgements

We would like to thank Matthias Landgraf, Eva Higginbotham, and Gabriel Aughey for feedback and advice on this project. We would like to thank Matthias Landgraf, Holly Ironfield, Benjamin White, Andrea Brand, Fernando J. Díaz-Benjumea, David John Luginbuhl, for providing fly stocks. For other fly stocks, we also thank Bloomington *Drosophila* Stock Center (NIH P40OD018537), the Vienna *Drosophila* Resource Center (VDRC, www.vdrc.at), and the FlyORF, Zurich ORFeome Project (https://flyorf.ch/). This work was funded by Wellcome Trust Investigator grant 104567 to T.D.S.

