## Supplemental figures for "Dynamic neurotransmitter specific transcription factor expression profiles during *Drosophila* development"

### Supplementary information

**Contents:**

**Figure S1.** RNA Pol II occupancy at neurotransmitter synthesis genes across development.

**Figure S2.** Altered chromatin accessibility at transcription factor and neurotransmitter synthesis genes.

**Figure S3.** *Dll* is enriched in cholinergic neurons in L3 larvae and switches to GABAergic enriched in adults.

**Figure S4.** *acj6* is expressed in cholinergic third instar larval neurons.

**Figure S5.** Absence of *apterous* expression in GABAergic third instar larval neurons.

**Figure S6.** Absence of *Ets65A-RA/C/D/E* expression in cholinergic third instar larval neurons.

**Figure S7.** Misexpression of candidate transcription factors are unable to alter neurotransmitter identity.

**Figure S8.** Neurotransmitter specific expression of non-coding RNAs during neural development.

**Table S1.** RNA Pol II bound genes (separate excel file)

**Table S2.** Neurotransmitter subtype enriched genes (separate excel file)

**Table S3.** Uniquely enriched and depleted transcription factors (separate excel file)

**Table S4.** Acj6 and Ets65A-PA bound genes (separate excel file)

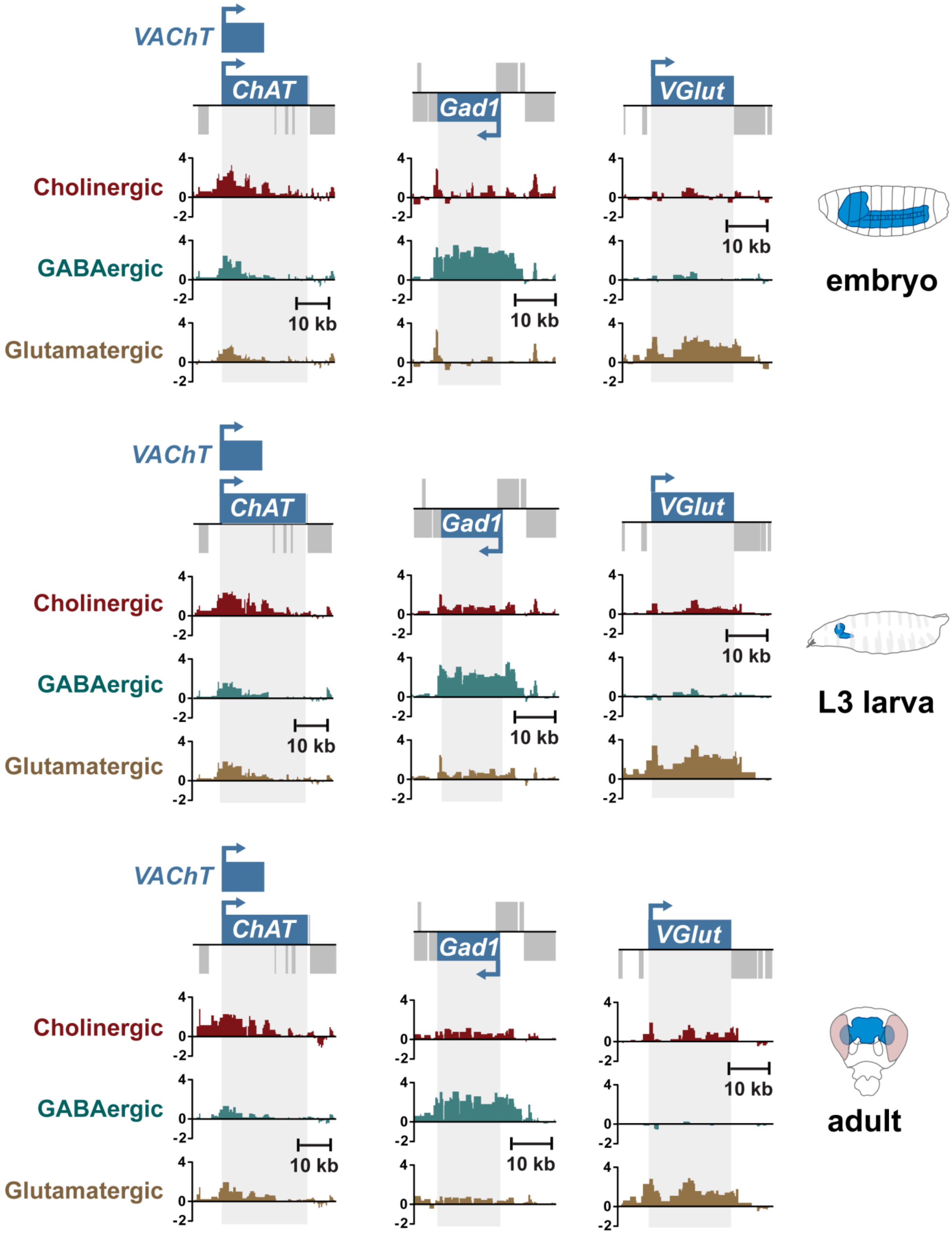


**Figure S1. RNA Pol II occupancy at neurotransmitter synthesis genes across development.**

Y-axis represent log2 ratios of Dam-Pol II over Dam-only.

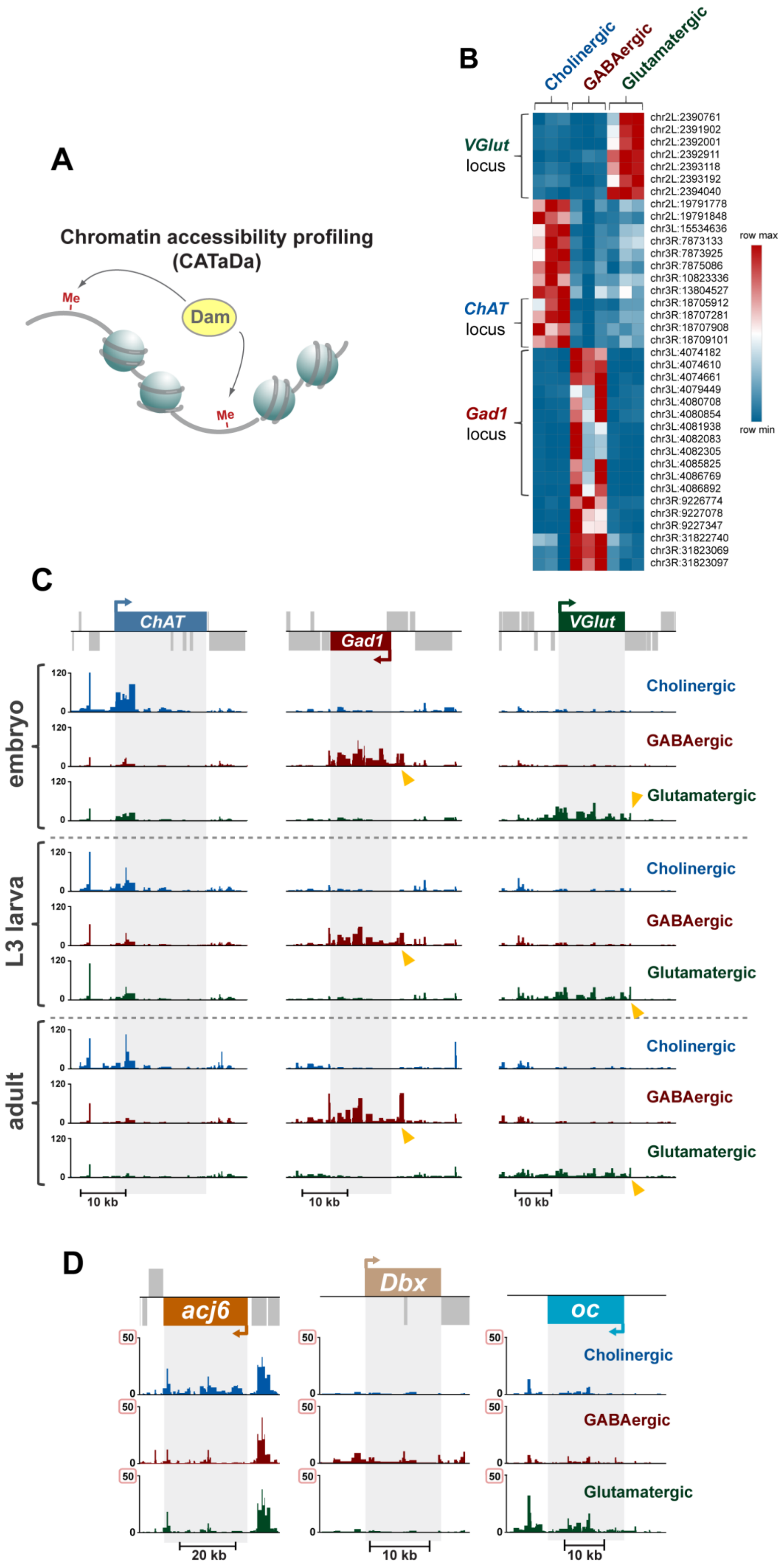


**Figure S2. Altered chromatin accessibility at transcription factor and neurotransmitter synthesis genes.** A) Chromatin Accessibility Targeted DamID (CATaDa) relies on methylation of open chromatin by Dam. B) Heatmap of GATC fragments with robust differences in accessibility (embryonic stage). C) Robust differences in accessibility at key neurotransmitter genes across all developmental stages. Y-axis represents RPM. Yellow arrows highlight accessible regions outside of the gene body. D) Minor differences (note the Y-axis maximum of 50 RPM) in accessibility (embryonic stage) at the *acj6*, *Dbx* and *oc* loci. Y-axis represents reads per million (RPM).

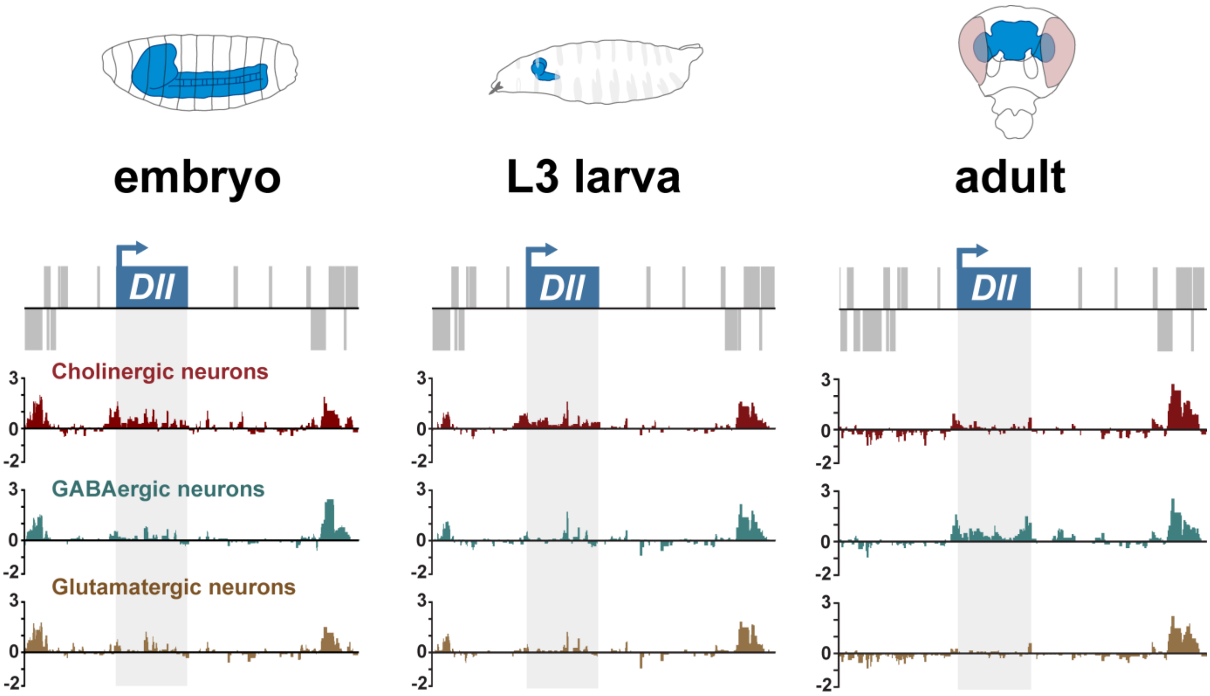


**Figure S3. *Dll* is enriched in cholinergic neurons in L3 larvae and switches to GABAergic enriched in adults.** Y-axis represent log2 ratios of Dam-Pol II over Dam-only.

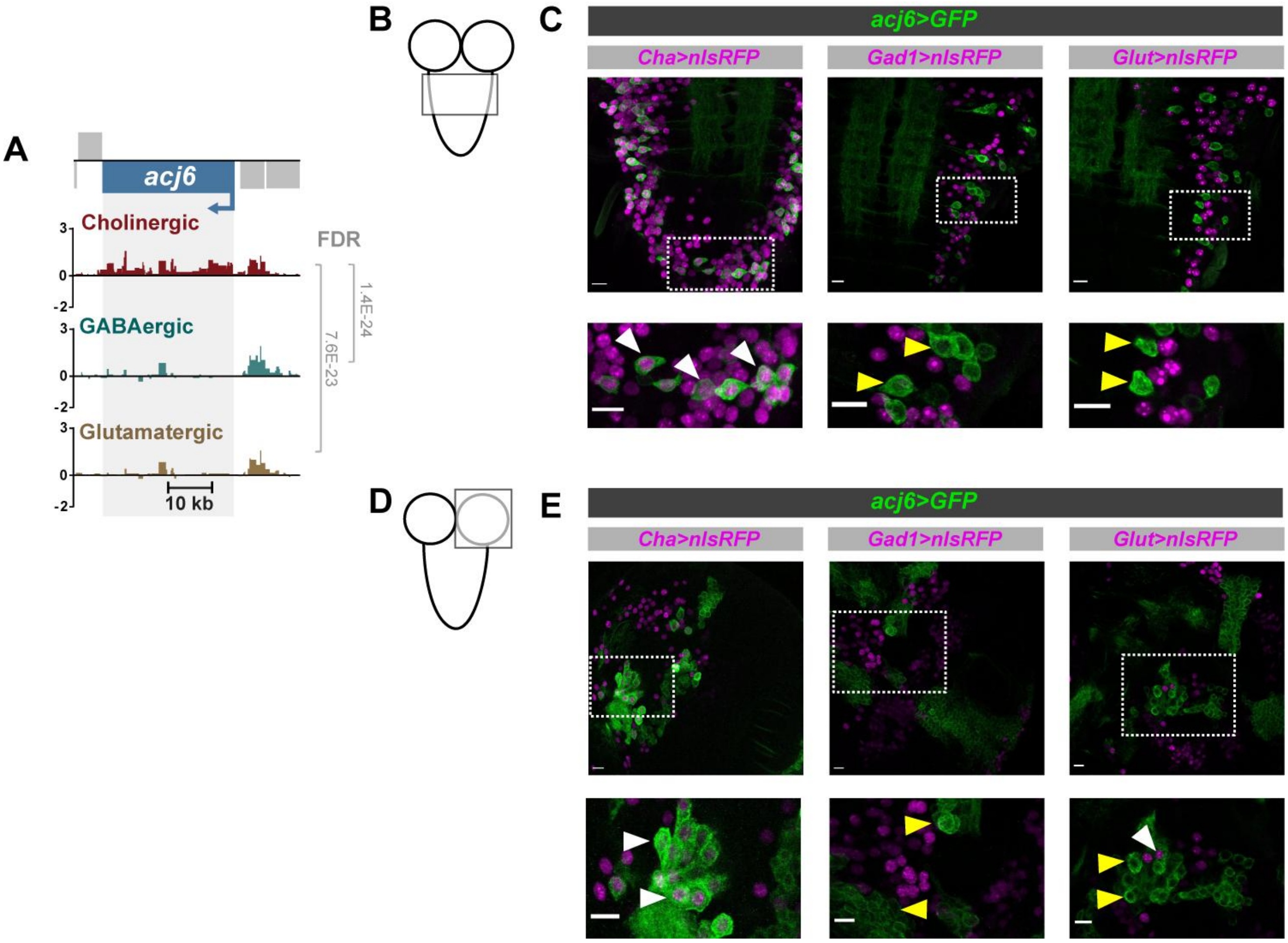


**Figure S4. *acj6* is expressed in cholinergic third instar larval neurons.** A) Pol II occupancy at *acj6* in third instar larval brain. Y-axis represents log2 ratio of Dam-Pol II over Dam-only. B) Drawing depicting a third instar laval CNS, the thoracic segments (imaged in C)) are highlighted with a rectangle. C) Confocal images of the Ventral Nerve Cord. D) Drawing depicting a third instar larval CNS. The right brain lobe (imaged in E)) is highlighted with a rectangle. E) Confocal images of right brain lobes. Outlined white rectangles show the zoomed area below. White triangles indicate coexpression, while yellow ones do not. All scale bars are 10 µm.

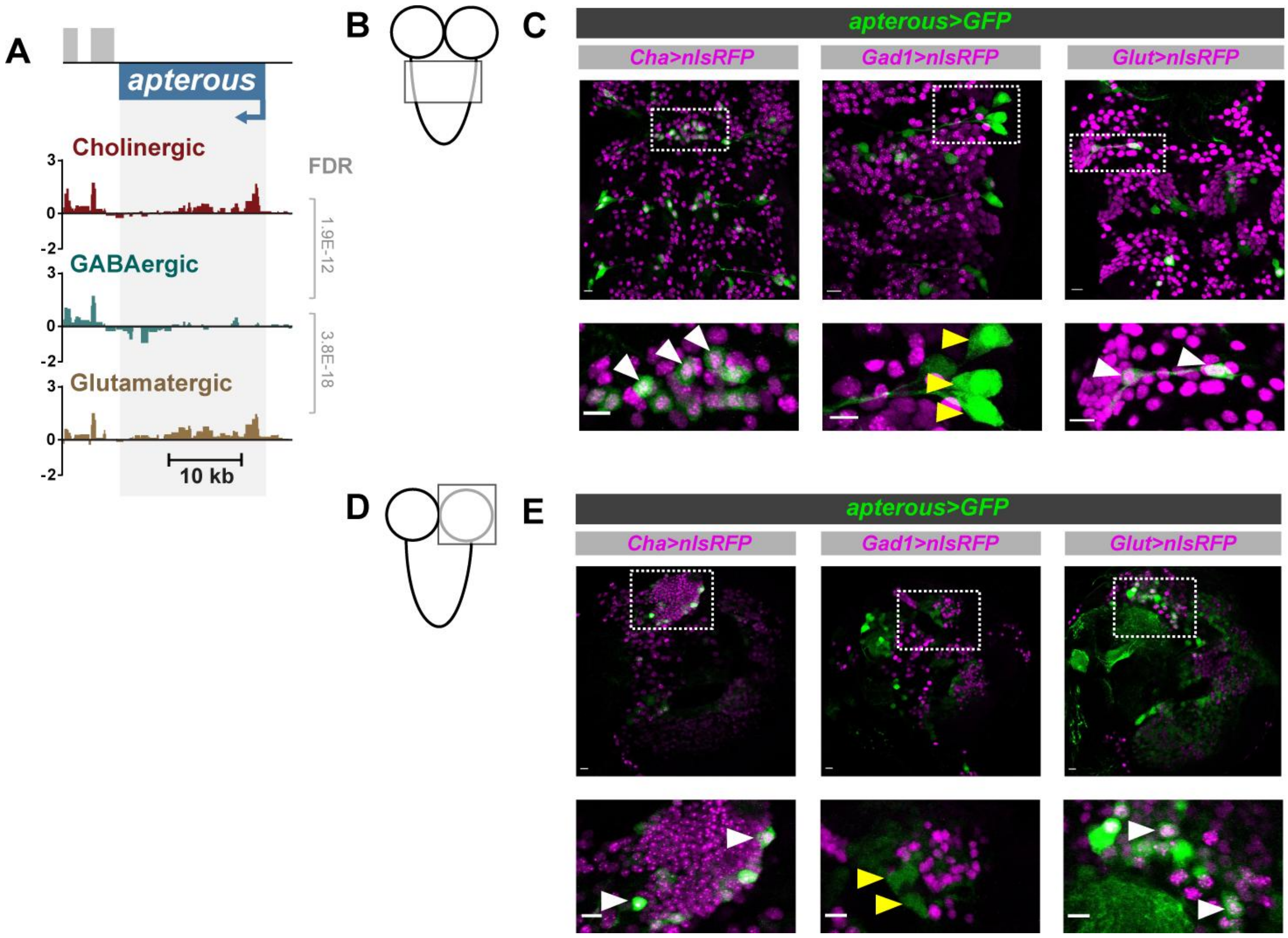


**Figure S5. Absence of *apterous* expression in GABAergic third instar larval neurons.** A) Pol II occupancy at *apterous* in third instar larval brain. Y-axis represents log2 ratio of Dam-Pol II over Dam-only. B) Drawing depicting a third instar laval CNS, the thoracic segments (imaged in C)) are highlighted with a rectangle.  C) Confocal images showing thoracic segments. D) Drawing depicting a third instar larval CNS. The right brain lobe (imaged in E)) is highlighted with a rectangle.  E) Confocal images showing right brain lobes. Outlined white rectangles showed the zoomed area below. White triangles indicate coexpression, and yellow ones do not. All scale bars are 10 µm.

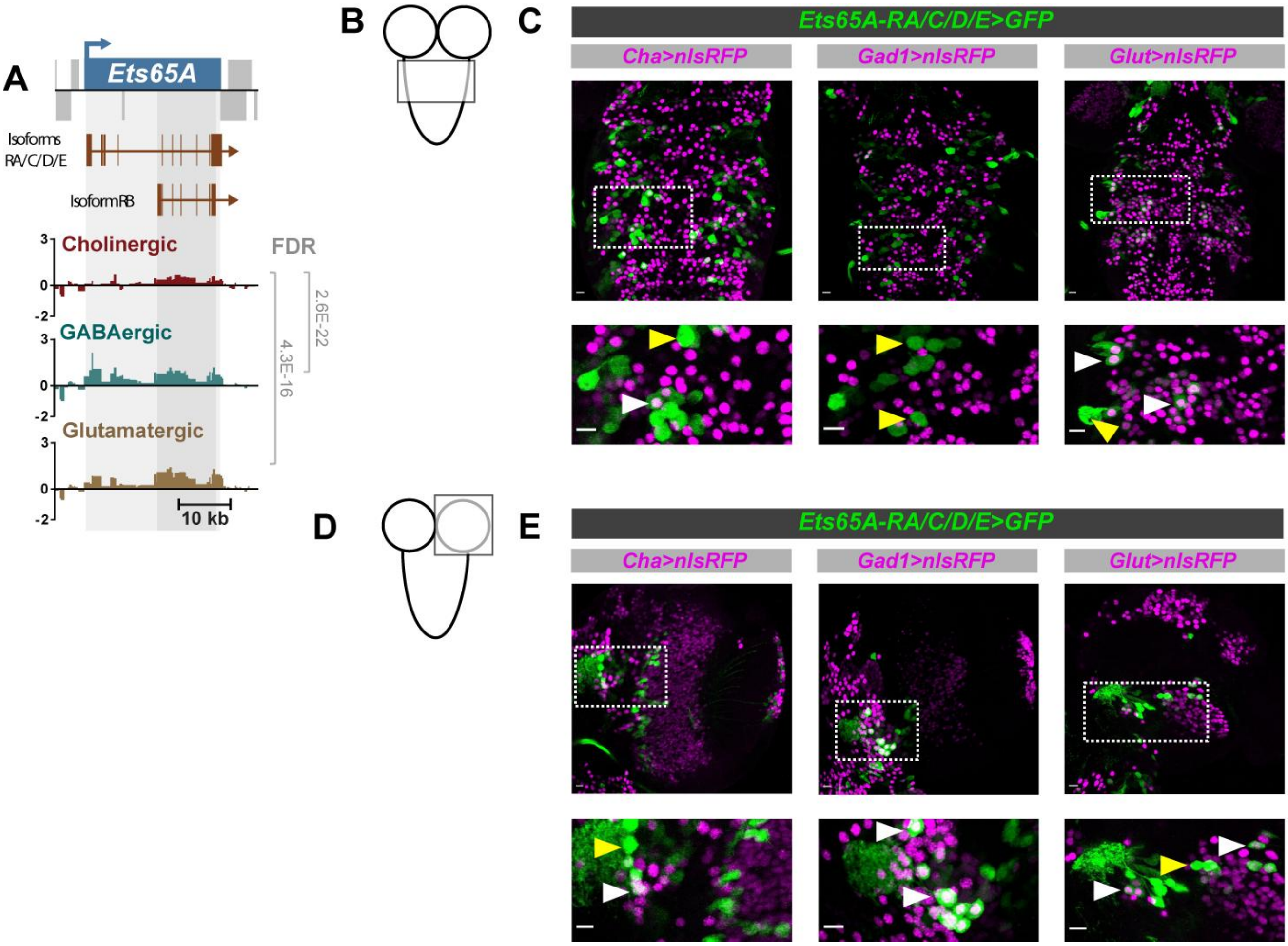


**Figure S6. Absence of *Ets65A-RA/C/D/E* expression in cholinergic third instar larval neurons.** A) Pol II occupancy at *Ets65A* in the third instar brain. Y-axis represents log2 ratio of Dam-Pol II over Dam-only. B) Drawing depicting a third instar laval CNS, the thoracic segments (imaged in C)) are highlighted with a rectangle. C) Confocal images showing thoracic segments. D) Drawing depicting a third instar larval CNS. The right brain lobe (imaged in E)) is highlighted with a rectangle. E) Confocal images showing right brain lobes. Outlined white rectangles show the zoomed area below. White triangles indicate coexpression and yellow ones do not. All scale bars are 10 µm.

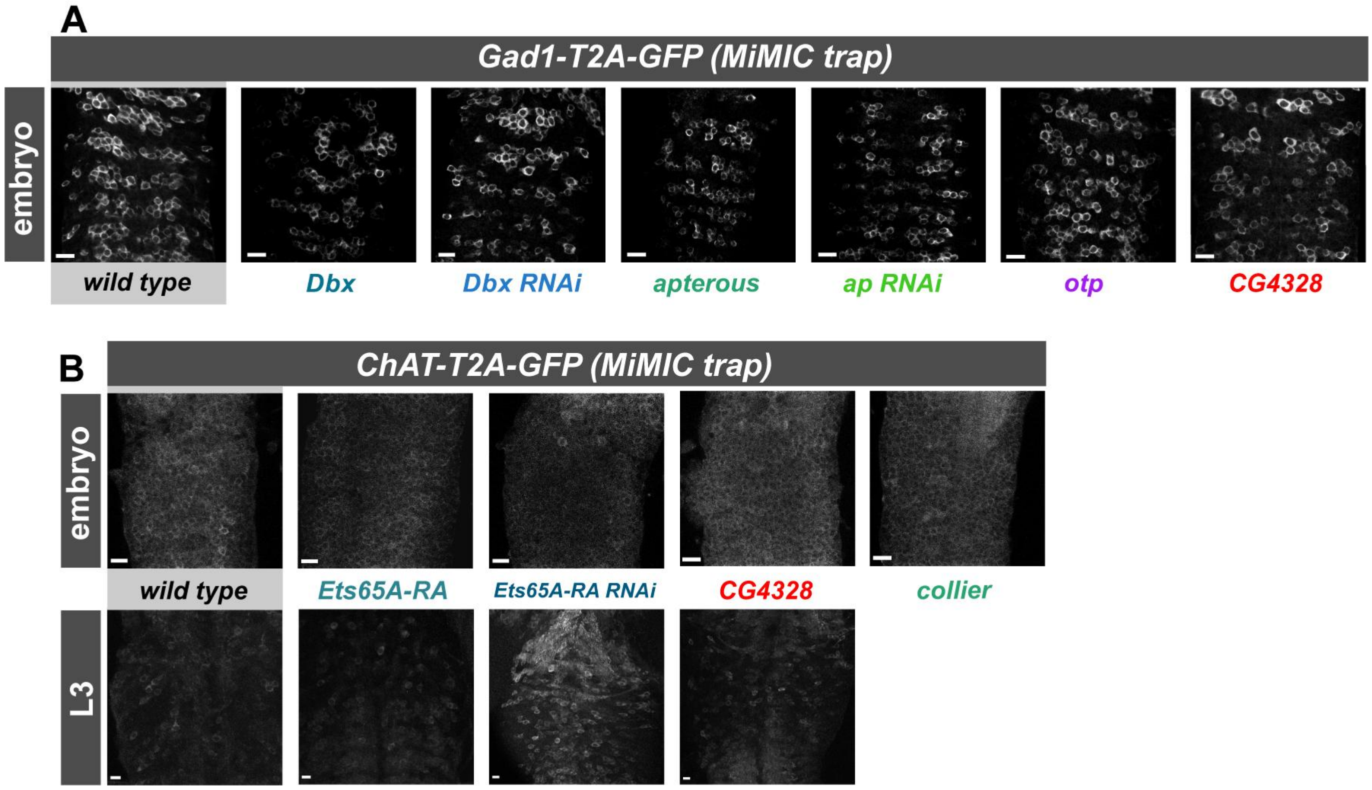


**Figure S7. Misexpression of candidate transcription factors are unable to alter neurotransmitter identity.** A) Gain of function and knockdown of candidates predicted to regulate GABAergic fate. Ventral images of stage 17 embryos. B) Gain of function and knockdown of candidates predicted to regulate cholinergic fate. Ventral images of stage 17 embryos, and thoracic segments of third instar larvae CNS. All scale bars represent 10 μm.

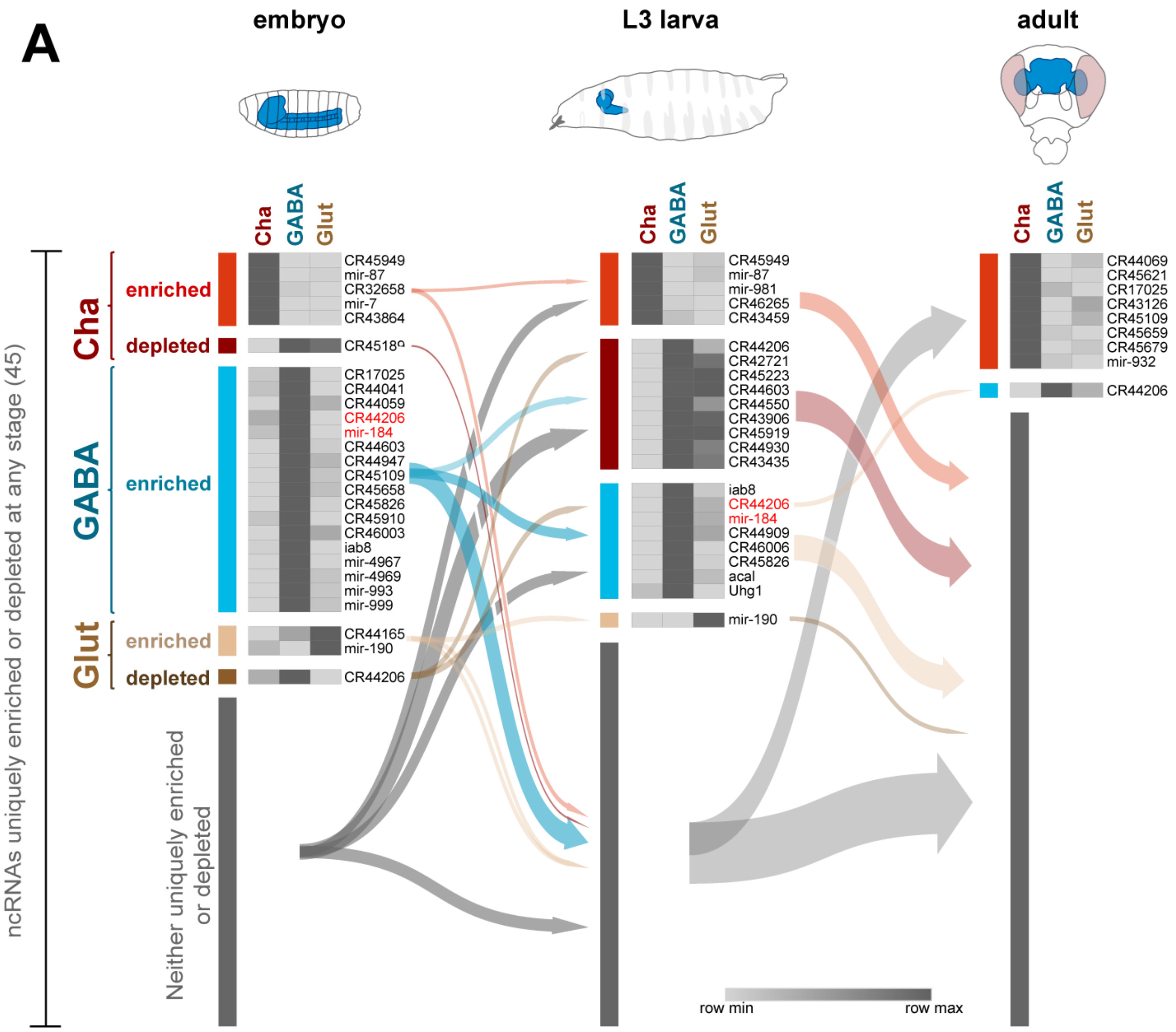


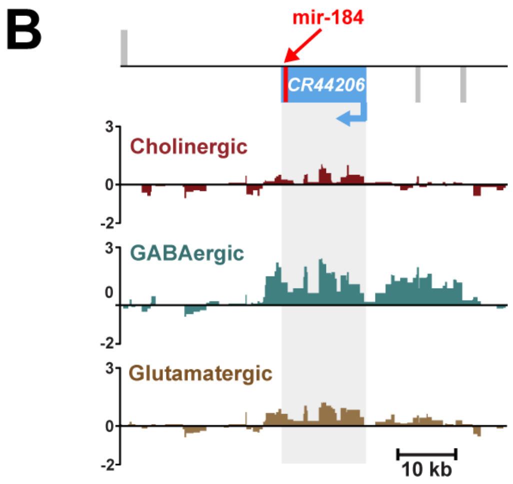


**Figure S8. Neurotransmitter specific expression of non-coding RNAs during neural development.** A) Non-coding RNAs uniquely enriched or depleted in cholinergic, GABAergic and glutamatergic neurons. A total of 45 transcription factors are identified across all stages. B) Enriched Pol II occupancy in GABAergic neurons at *mir-184* and *CR44206* in L3 larval brains.
